# Gut microbiota profiles of treatment-naïve adult acute myeloid leukemia patients with neutropenic fever during intensive chemotherapy

**DOI:** 10.1101/2020.07.09.194910

**Authors:** Thanawat Rattanathammethee, Pimchanok Tuitemwong, Parameth Thiennimitr, Phinitphong Sarichai, Sarisa Na Pombejra, Pokpong Piriyakhuntorn, Sasinee Hantrakool, Chatree Chai-Adisaksopha, Ekarat Rattarittamrong, Adisak Tantiworawit, Lalita Norasetthada

## Abstract

Intestinal bacterial flora of febrile neutropenic patients had significantly diverse. However, there were scanty reports of microbiota alteration of adult patients with acute myeloid leukemia (AML). Stool samples of each treatment-naïve AML patient were collected at the day before induction chemotherapy initiation (pretreatment), first day of neutropenic fever and first day of bone marrow recovery. Bacterial DNA was extracted from stool and sequenced bacterial 16s ribosomal RNA genes by next-generation sequencing. Relative abundance, overall richness, Shannon’s diversity index and Simpson’s diversity index were calculated. Ten cases of AML patients (4 men and 6 women) were included with median age of 39 years (range: 19-49) and all of patients developed febrile neutropenia. Firmicutes were dominated over the period of neutropenic fever and subsequent declined after bone marrow recovery contrast to Bacteroidetes and Proteobacteria. *Enterococcus* was more abundant at febrile neutropenia period compared to pretreatment while *Bacteroides* and *Escherichia* was notably declined during the febrile neutropenia. At the operational taxonomic units (OTUs) level, there was significant higher level of overall richness of pretreatment period than febrile neutropenic episode. Both of the diversity indexes of Shannon and Simpson were considerably decreased at febrile neutropenic period. Adult AML patients with first episode of febrile neutropenia after initial intensive chemotherapy demonstrated the significant decrease of gut microbiota diversity and the level of diversity consistently remained constant despite of bone marrow recovery.

## Introduction

Up to half of patients with solid tumors and over 80% of those with hematologic malignancies develop a fever during chemotherapy-induced neutropenia.[1] Recommended current clinical practice includes broad-spectrum antibiotics at the onset of neutropenic fever (NF), while most of NF patients remain negative microbiological workup.[2]This practice of empiric antimicrobial attack rather than a mechanistic approach by precisely defined pathogenesis. Moreover, this approach has led to serious adverse consequences, including antibiotic resistance and *Clostridium difficile* infection.

Neutropenic fever after intensive chemotherapy associated to iatrogenic damage to the gut microbiota. Intensive chemotherapy has also impaired gut barrier integrity, facilitating bacterial translocation and lead to increasing risk of bloodstream infection.[3] Additionally, there were evidences of gut microbiota disruption that normally prevent pathogen colonization[4], provide tonic stimulation to gut barrier[5] and facilitate recovery from chemotherapy-induced injury after empirical antibiotics in NF patients.[6]

Although the majority of previous gut microbiota studies [7–12] were based on patients who received stem cell transplantation that duration of neutropenic period particularly longer than the other kinds of chemotherapeutic categories, gastrointestinal bacterial colonization will often be affected during first treatment course of acute leukemia both through mucosal barrier injuries, the use of broad spectra antibiotics and other antimicrobial agents.[13]

This study aims to explore gut microbiota profiles in patients with NF during intensive chemotherapy that further information could reveal gut dysbiosis, an imbalanced gut microbiota, and specify the timing and other specifics antibiotic de-escalation. We designated to compare the alteration of gut microbiota at pretreatment, during neutropenic fever and recovery phase of neutropenia. Thus, we sought to test the hypothesis that gut microbiota composition could be altered in patients with acute myeloid leukemia (AML) who developed first episode of neutropenic fever during first cycle of intensive chemotherapy.

## Materials and methods

### Study population

We enrolled ten consecutive Thai patients of treatment-naïve AML undergoing first cycle of induction chemotherapy between July to September 2017 in Hematology Unit of Maharaj Nakorn Chiang Mai Hospital, Chiang Mai University, Thailand. All of included patients aged 18 to 65 years and overall condition suited to intensive treatment. The diagnosis of AML defined as the presence of blasts of myeloid series more than 20% in circulation and/or bone marrow examination according to WHO classification of myeloid neoplasm.[14] All patients received standard induction chemotherapy “7+3 regimen” (seven-day of Cytarabine 100 mg/m^2^ intravenous continuous infusion over 24 hours combine to three-day of Idarubicin 12 mg/m^2^ bolus intravenously). All patients developed NF which defined as single oral temperature of ≥ 101°F (38.3°C) or a temperature of ≥ 100.4°F (38°C) sustained over 1 hour plus an absolute neutrophil count (ANC) of < 0.5 × 10^9^/L.[2] Single agent Piperacillin/Tazobactam was first empirical antibiotic for NF patients and subsequent treatment was allowed regarding to patient’s condition and physician’s decision according to international guideline. [2] Granulocyte-colony stimulating factor (G-CSF) was not allowed to administer in all participants.

We excluded patients who previously treated with antibiotics within 90 days and/or probiotics and patients who received nasal tube feeding or parenteral nutrition during the study period, as these factors are well described as impacting the intestinal microbiota. [15, 16] The fluoroquinolones prophylaxis was not used in our center, but all of patients received prophylactic dose of itraconazole (200 mg twice daily) and acyclovir (400 mg twice daily) for aspergillosis and herpes zoster reactivation, respectively.

The institutional ethical review board of Faculty of Medicine, Chiang Mai University, Thailand, approved the study. Written informed consent was obtained from all the participants before they accepted to enroll in the study. (study code: MED-2559-03947)

### Sample collection

The stool samples were collected from 10 AML patients. Stool was collected by standard stool kit including sterile plastic cup with lid and plastic bag with zip lock to cover all of specimens. Fecal samples were stored at −20 °C prior to DNA extraction. Three episodes of stool sampling were indicated; pretreatment (at the day before starting of chemotherapy), the first day of febrile neutropenia and first day of bone marrow recovery implied by surge of ANC more than 0.5 × 10^9^/L for 24 to 48 hours apart in two consecutive times without any transfusion supported for maintain the appropriated level of red blood cells (more than 7 g/dl of hemoglobin [Hb] level) and platelet count (above of 10 × 10^9^/L without bleeding symptoms).[17] All participants were controlled of dietary consumption for neutropenic patients according to hospital’s dietary policy. All the samples were collected by individual patient. (Fig 1)

**Fig 1.**
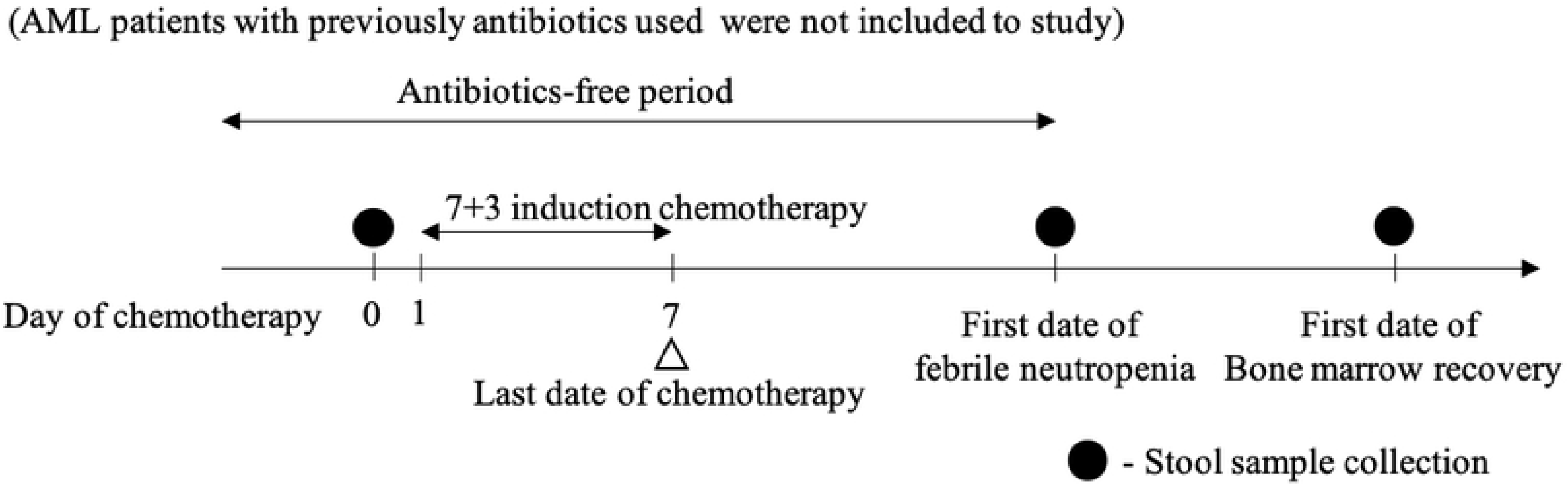
Study schema. Stool sample collection were collected from 10 consecutive adults with acute myeloid leukemia receiving 7+3 induction chemotherapy at pretreatment, first date of febrile neutropenia and first date of bone marrow recovery.

### Bacterial stool DNA extraction

DNA extraction from AML patients’s stool sample was performed using QIAamp DNA Stool Mini Kit (Qiagen, Hilden, Germany) according to the manufacturer’s instructions. The obtained DNA was quantified using a spectrophotometer (NanoDrop Technologies, Wilmington, DE).

### Polymerase chain reaction and sequencing

The DNA samples were sent to Omics sciences and bioinformatics center of Chulalongkorn University (Bangkok, Thailand) to perform the next generation sequencing (NGS) analysis. Polymerase chain reaction (PCR) amplification of the V3-V4 region of the bacterial 16s ribosomal RNA (16s rRNA) genes was performed using broad spectrum 16s rRNA primers [18] (forward primer: 5’-TCGTCGGCAGCGTCAGATGTGTATAAGAGACAGCCTACG GGNGGCWGCAG-3’ and reverse primer 5’-GTCTCGTGGGCTCGGAGATGTGTATAAG AGACAGGACTACHV GGGTATCTAATCC-3’). Amplicons were generated using a high-fidelity polymerase, 2X KAPA hot-start ready mix (KAPA Biosystems, USA). The amplification condition included an initial denaturation step 3 minutes at 94 °C, followed by 25 cycles of 98 °C for 20 seconds, 55 °C for 30 seconds, and 72 °C for 30 seconds, followed by a single step final extension step at 72 °C for 5 minutes. The targeted amplicons were purified using a magnetic bead capture kit (Agencourt AMPure XP, Beckman Coulter, USA). Subsequently, the purified 16S amplicons were indexed using 2X KAPA hot-start ready mix and 5 μl of each Nextera XT index primer in a 50 μl PCR reaction, followed by 8-10 cycles of PCR condition as above, purified using AMPure XP beads, pooled and diluted to final loading concentration at 6 pM. Sequencing was performed using the Illumina 16s MiSeq sequencing system, according to standard operating procedures, with read length 250 bases in paired-end sequencing mode.

### Sequencing analysis

Sequencing reads quality examined using FASTQC software. [19] Overlapping paired end reads were assembled using PEAR. FASTX-Toolkit is used to filter out assembled reads that do not have a quality score of 30 at least 90% of bases, and then remove reads that are less than 400 base-pair long. Chimeras were removed by the UCHIME method [20] as implemented in vsearch1.1.1 [21] using –uchime_ref option against chimera-free Gold RDP database. Using the *pick_open_reference_otus.py* command in Quantitative Insights Into Microbial Ecology (QIIME) 1.9.0 pipeline [22] to determine the operational taxonomic units (OTUs) which corresponded to the 16s rRNA gene sequences to address the microbial diversity and using BLAST analysis [23] for non-redundant 16s rRNA reference sequences, which were obtained from the Ribosomal Database Project. [24] Taxonomic assignment was based on NCBI Taxonomy. [25]

### Statistical analysis

All results of 16s rRNA gene sequencing were assigned to each category of bacteria as phylum to genus level. The data were entered into custom database (Excel, Microsoft Corp) and analyzed using Prism 8 software (GraphPad, Inc., La Jolla, CA). Quantitative data were reported as mean ± SD or median (range). The relative abundance, the overall richness by comparison of OTUs, and the Shannon [26] and Simpson [27] diversity indexes at the phylum level were calculated and statistically performed using one-way analysis of variance (alternatively, the Kruskal-Wallis test) to compare each clinical timepoints of individual patient and paired t-test (alternatively, the Wilcoxon signed-rank test) were used to compare paired samples. Statistical corrections for multiple comparisons were performed using Bonferroni methods.

## Results

### Patients’ characteristics

Ten cases of AML (4 men and 6 women) were included with median age of 39 years (range: 19-49 years). All of patients received 7+3 induction regimen and developed NF, initial empirical antibiotics was Piperacillin/Tazobactam (100%) and adjustment of antibiotics and antifungal was allowed regarding to clinical course of individual patient by physician’s decision (Fig 2). Three patients (33.3%) were microbiological defined. All cases had invasive pulmonary aspergillosis confirmed by typical radiologic finding and serum galactomannan (indicated as patient’s code: P4, P6 and P10) and two of them were co-infected with *Pseudomonas* pneumonia (P6) and *Escherichia coli* septicemia (P10). The gastrointestinal symptoms during hospitalization included nausea (100%) and watery diarrhea (10%, P10). (Table 1) Median ANC were 2.85 × 10^9^/L (range: 1.42-7.67 × 10^9^/L), 0.04 × 10^9^/L (range: 0.01-0.43 × 10^9^/L) and 3.65 × 10^9^/L (range: 2.09-5.78 × 10^9^/L) at the day before treatment initiation, first date of febrile neutropenia and first date of bone marrow recovery, respectively. Median time to neutropenia was 11 days (range: 8-13 days) and median duration of neutropenia was 12 days (range: 7-17 days). Days of administration of antibiotics, antifungal agents and stool sample collection of individual patients were presented in figure 2. In total, 24 stool samples were collected from 10 AML patients. The samples were assigned to three groups: (1) Pretreatment (n = 10); (2) Febrile neutropenia (n=9); and (3) Bone marrow recovery (n=5). All of the missing stool samples were resulted from technical issues.

**Fig 2.**
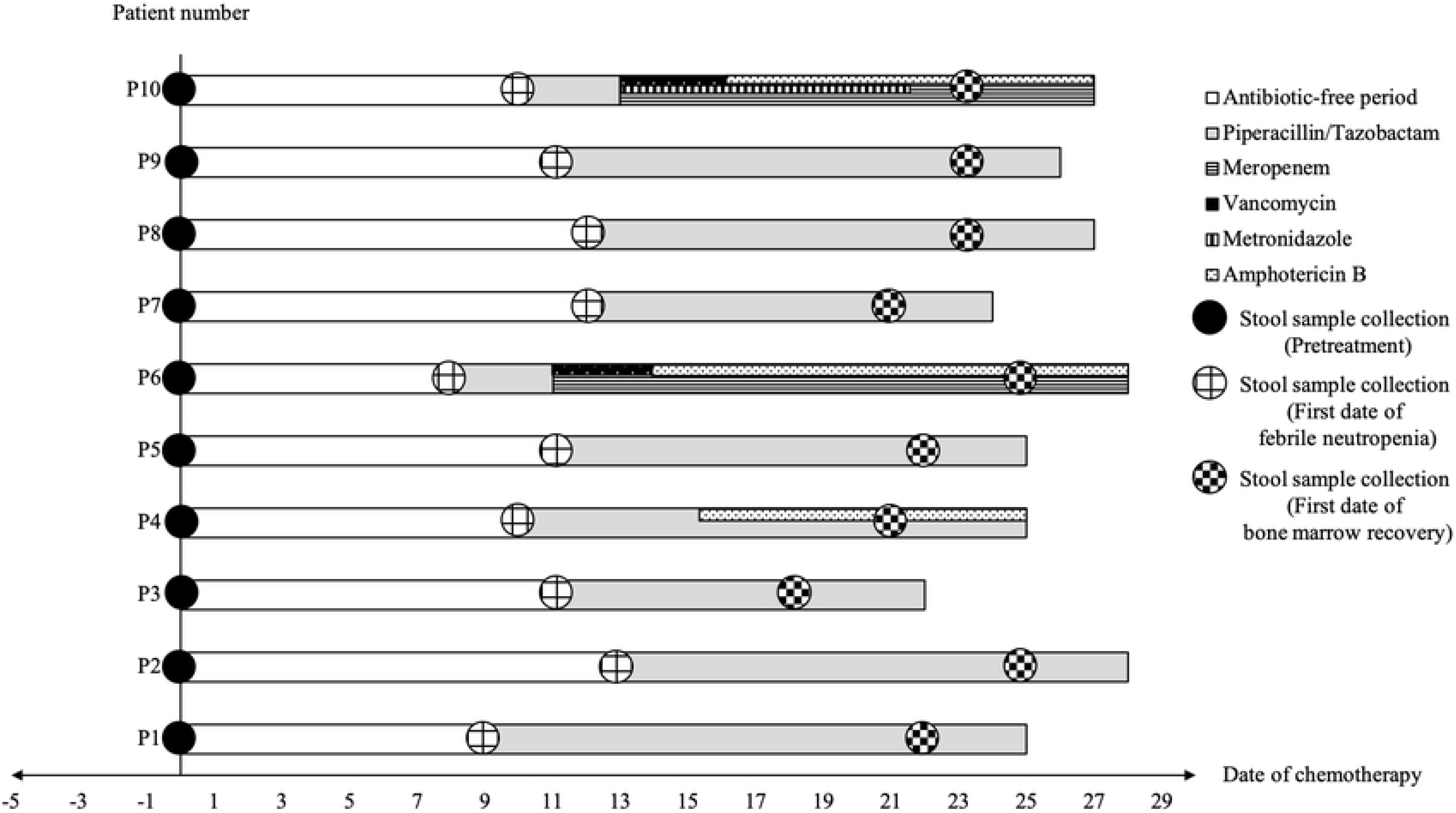
Antibiotics, antifungal and stool sample collection of each individual patients.

**Table 1.** Characteristics of the ten AML patients included in the study.

| Code | Sex | Age | Time to neutropenia (days) | Neutropenia duration (days) | Microbiological defined events | ANC ( x 10<sup>9</sup>/L) |  |  |
| --- | --- | --- | --- | --- | --- | --- | --- | --- |
|  |  |  |  |  |  | Pre-treatment | FN | BM recovery |
| P1 | Male | 41 | 9 | 13 | none | 2.49 | 0.01 | 3.01 |
| P2 | Female | 37 | 13 | 12 | none | 3.93 | 0.03 | 5.78 |
| P3 | Male | 35 | 11 | 7 | none | 1.42 | 0.16 | 3.43 |
| P4 | Female | 24 | 10 | 11 | IPA | 7.67 | 0.08 | 5.03 |
| P5 | Male | 40 | 11 | 11 | none | 5.36 | 0.05 | 3.62 |
| P6 | Female | 19 | 8 | 17 | *Pseudomonas pneumonia*; IPA | 2.35 | 0.01 | 4.54 |
| P7 | Male | 44 | 12 | 9 | none | 1.87 | 0.02 | 3.52 |
| P8 | Female | 49 | 12 | 12 | none | 3.22 | 0.25 | 2.09 |
| P9 | Female | 44 | 11 | 13 | none | 2.38 | 0.02 | 3.68 |
| P10 | Female | 20 | 10 | 14 | *E.coli* septicemia; IPA | 3.70 | 0.43 | 3.89 |
| Median (range) |  | 39 (19-49) | 11 (8-13) | 12 (7-17) | - | 2.85 (1.42-7.67) | 0.04 (0.01-0.43) | 3.65 (2.09-5.78) |

ANC, Absolute neutrophil count; FN, Febrile neutropenia; BM, Bone marrow; *E.coli*, *Escherichia coli*; IPA, Invasive pulmonary aspergillosis

### Distribution of bacterial phyla in the gut microbiota among AML patients

Figure 3 shows the relative abundance of the bacterial phyla in each timepoints of AML treatment. Across all the samples, the following five most abundant bacterial phyla were identified (Table 2): Firmicutes (41.7%), Bacteroidetes (28.7%), Proteobacteria (17.6%), Verrucomicrobia (7.8%) and Spirochaetes (2.3%). At the first date of febrile neutropenia, Firmicutes predominately rose from median of relative abundance of 34.3% to 50.8% and subsequent declined after bone marrow recovery. In contrast, Bacteroidetes and Proteobacteria were similarly dropped at NF period before level up at the last time of sample collection. Verrucomicrobia and Spirochates showed minimal alteration. However, there were no significant difference of relative abundance in each timepoint at phylum level.

**Fig 3.**
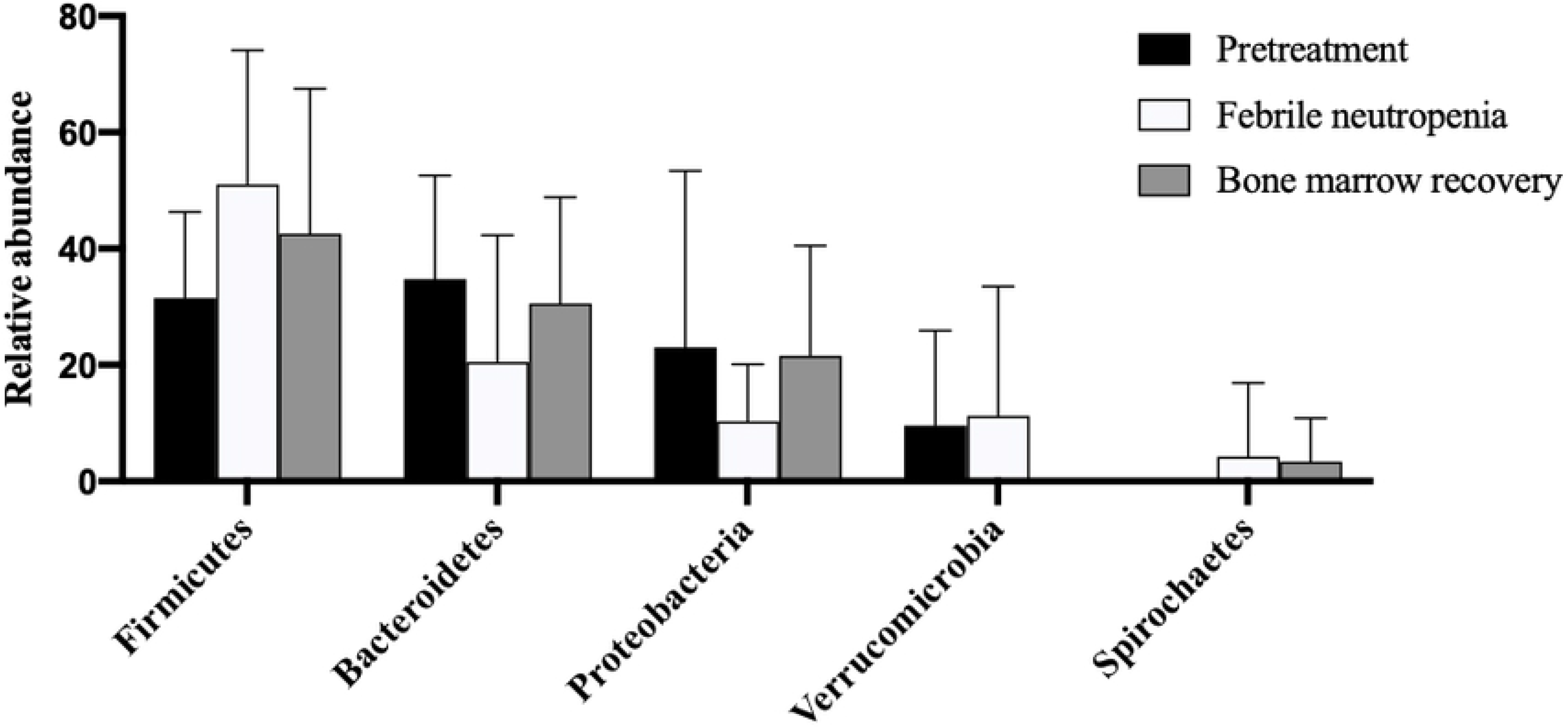
Relative abundance of the bacterial phyla in gut microbiota among ten cases of acute myeloid leukemia patients. Values shown are means ± standard deviation (SD).

**Table 2.**
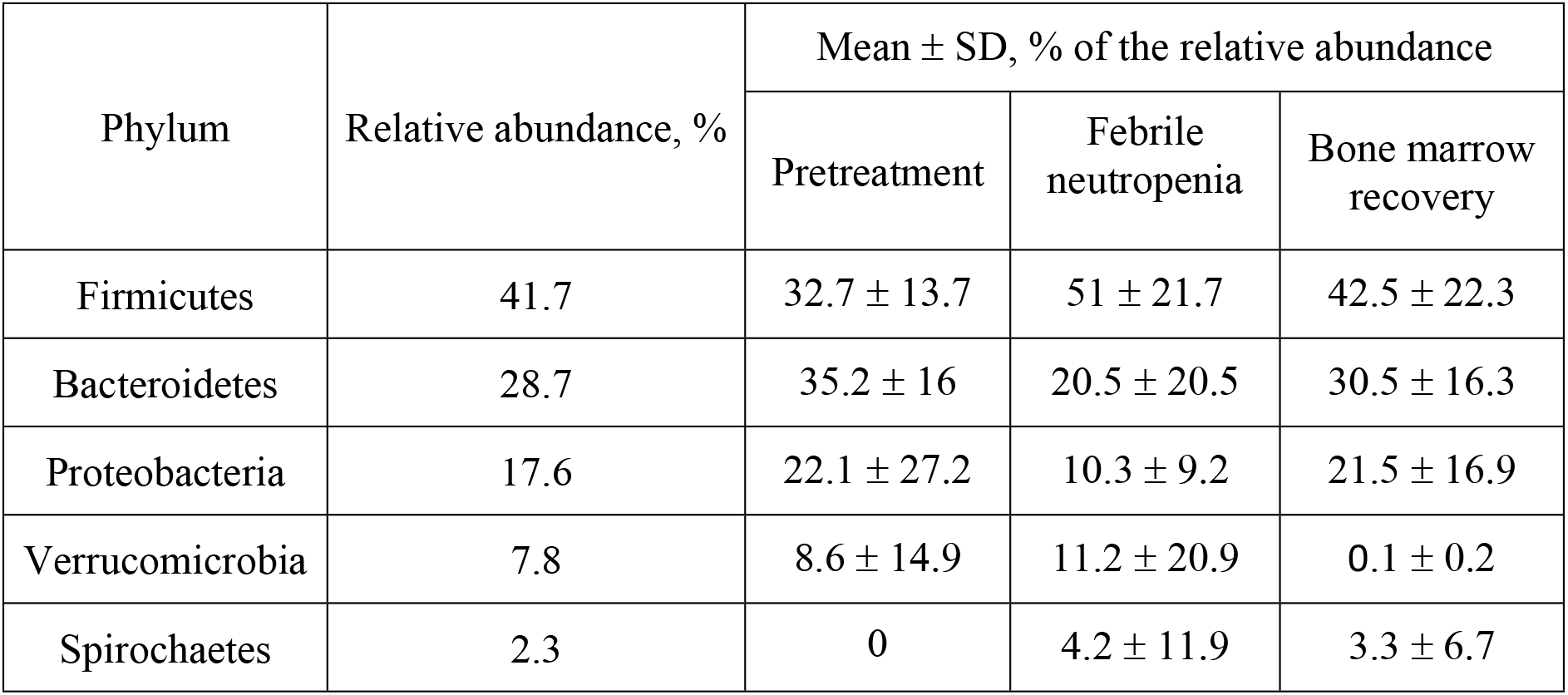
Relative abundance of the five most abundant bacterial phyla.

### Relative abundance at genus level

The genus extraction of phylum that relative abundance over 10% (Firmicutes, Bacteroidetes and Proteobacteria) were examined to determine specific bacterial organisms (Table 3). In phylum Firmicutes, the following bacterial genus were identified: *Enterococcus* (11.1%), *Blautia* (3.9%), *Streptococcus* (1.2%) and *Veillonella* (1.2%). *Enterococcus* was more abundant at febrile neutropenia period compared to pretreatment (mean difference 20.2, [95%CI (5.9, 34.6)]; *P* <0.01) and decrease over bone marrow recovery phase.

**Table 3.** Relative abundance at genus level of phylum that relative abundance over 10%.

| Phylum | Genus | Mean difference (95% Confidence interval) |  |  |
| --- | --- | --- | --- | --- |
|  |  | Pretreatment<br>vs.<br>Febrile neutropenia | Pretreatment<br>vs.<br>BM recovery | Febrile neutropenia<br>vs.<br>BM recovery |
| Firmicutes | *Enterococcus* | 20.2**<br>(5.9, 34.6) | 15.4<br>(-1.6, 32.5) | -4.7<br>(-22.2, 12.6) |
|  | *Streptococcus* | 0.7<br>(-13.6, 15.1) | 2.6<br>(-14.4, 19.7) | 1.9<br>(-15.4, 19.3) |
|  | *Blautia* | -1.8<br>(-16.2, 12.4) | -2.4<br>(-19.5, 14.6) | 0.6<br>(-18, 16.8) |
|  | *Veillonella* | -0.8<br>(-15.2, 13.4) | 2.1<br>(-15, 19.1) | 2.9<br>(-14.4, 20.3) |
| Bacteroidetes | *Bacteroides* | -11.7*<br>(-21.9, -1.4) | -6.7<br>(-26, 12.4) | 3.3<br>(-16, 22.8) |
|  | *Parabacteroides* | -1.5<br>(-17.1, 14.1) | 3.7<br>(-15.5, 22.9) | 5.2<br>(-14.1, 24.7) |
| Proteobacteria | *Sutterella* | -2.5<br>(-13.6, 8.6) | -2.8<br>(-16.4, 10.7) | -0.3<br>(-14.1, 13.4) |
|  | *Escherichia* | -11.6*<br>(-22.7, -0.4) | -9.1<br>(-22.7, 4.5) | 2.4<br>(-11.3, 16.2) |
|  | *Klebsiella* | 1.8<br>(-9.3, 12.9) | 4.7<br>(-8.8, 18.3) | 2.9<br>(-10.8, -16.7) |

BM: Bone marrow; \*\*p < 0.01; \*p < 0.05

Bacteroidetes phylum, the genus *Bacteroides* and *Parabacteroides* were extracted with relative abundance of 21.5% and 3.8%, respectively. In contrary to *Enterococcus*, *Bacteroides* was significantly decreased during the febrile neutropenia (mean difference - 11.7, [95%CI (−21.9, −1.4)]; *P* = 0.027) and upturn after bone marrow recovery.

For Proteobacteria phylum, *Sutterella*, *Escherichia* and *Klebsiella* were detected and accounted to 1.1%, 11.3% and 2.3%, respectively. Likewise, to *Bacteroides* at neutropenic fever, the *Escherichia* was notably declined (mean difference −11.6, [95%CI (−22.7, −0.4)]; *P* = 0.034) and rebound up after bone marrow recovery period.

### Richness and diversity of the gut microbiota among AML patients

To assess microbiota richness, the numbers of OTUs per patient were calculated (Fig 4). Pretreatment period showed a significant higher mean number of OTUs compared to febrile neutropenic episode (203.1 vs. 131.7; *P* = 0.012). However, there were no significant difference between OTUs of bone marrow recovery to pretreatment and first date of febrile neutropenia.

**Fig 4.**
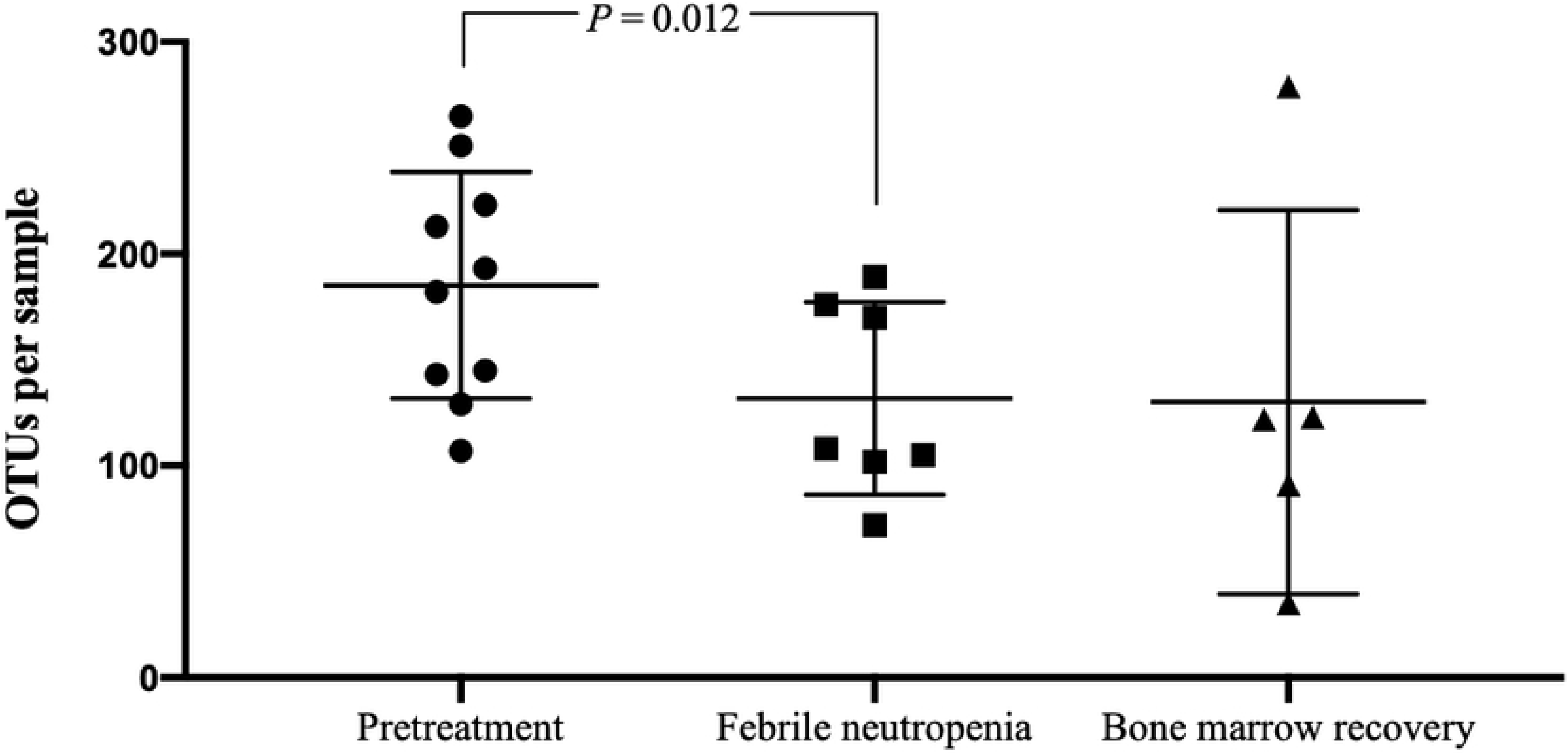
Richness of the gut microbiota in acute myeloid leukemia patients. Bacterial DNA was extracted and the 16s rRNA genes were sequenced and assigned to operational taxonomic units (OTUs). Each symbol represents one individual sample. Values shown are means ± SD.

The diversity indexes, Shannon and Simpson, were compared at the phylum level. (Figs 5a and 5b). It was found that the Shannon and Simpson diversity index showed significant reduction of bacterial abundance at febrile neutropenic period compare to pretreatment. (median of Shannon’s index of 1.077 vs. 1.002; *p* = 0.044, and median of Simpson’s index of 0.628 vs. 0.521; *p* = 0.027)

**Fig 5.**
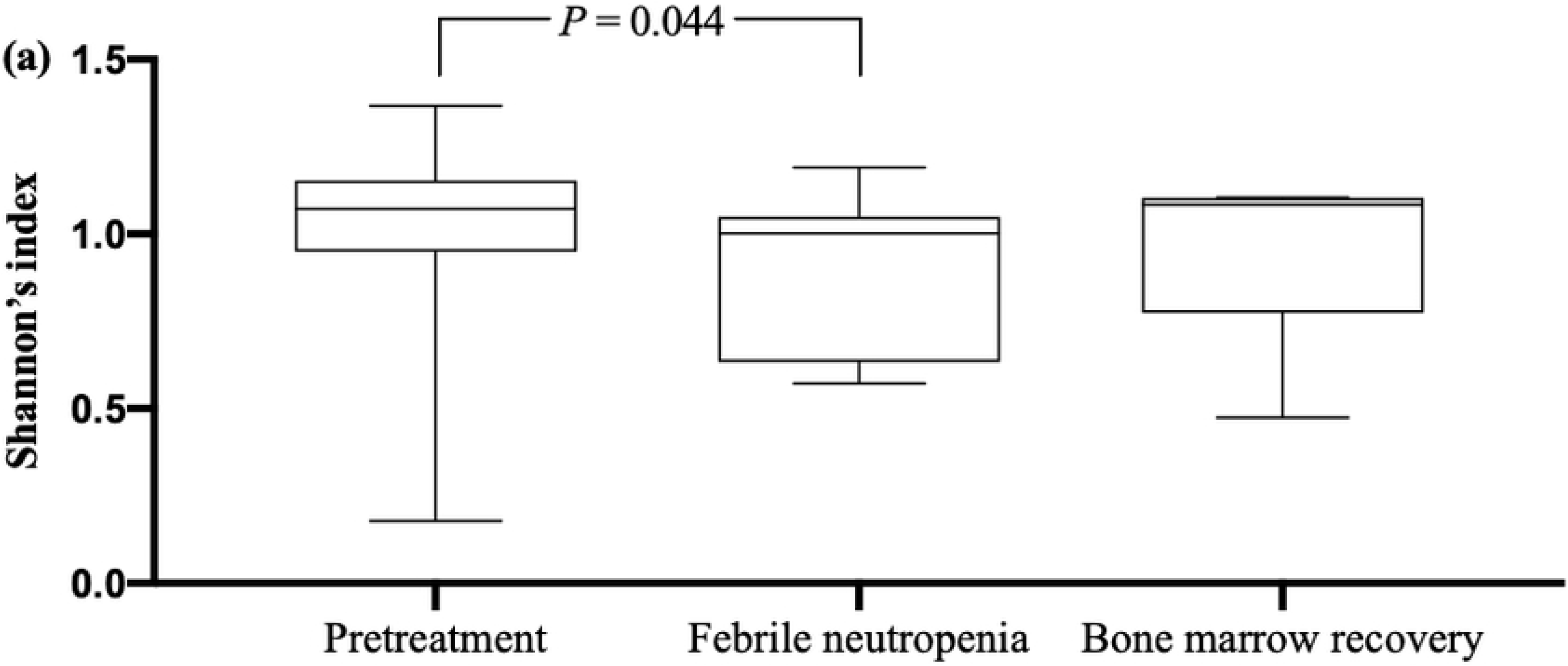
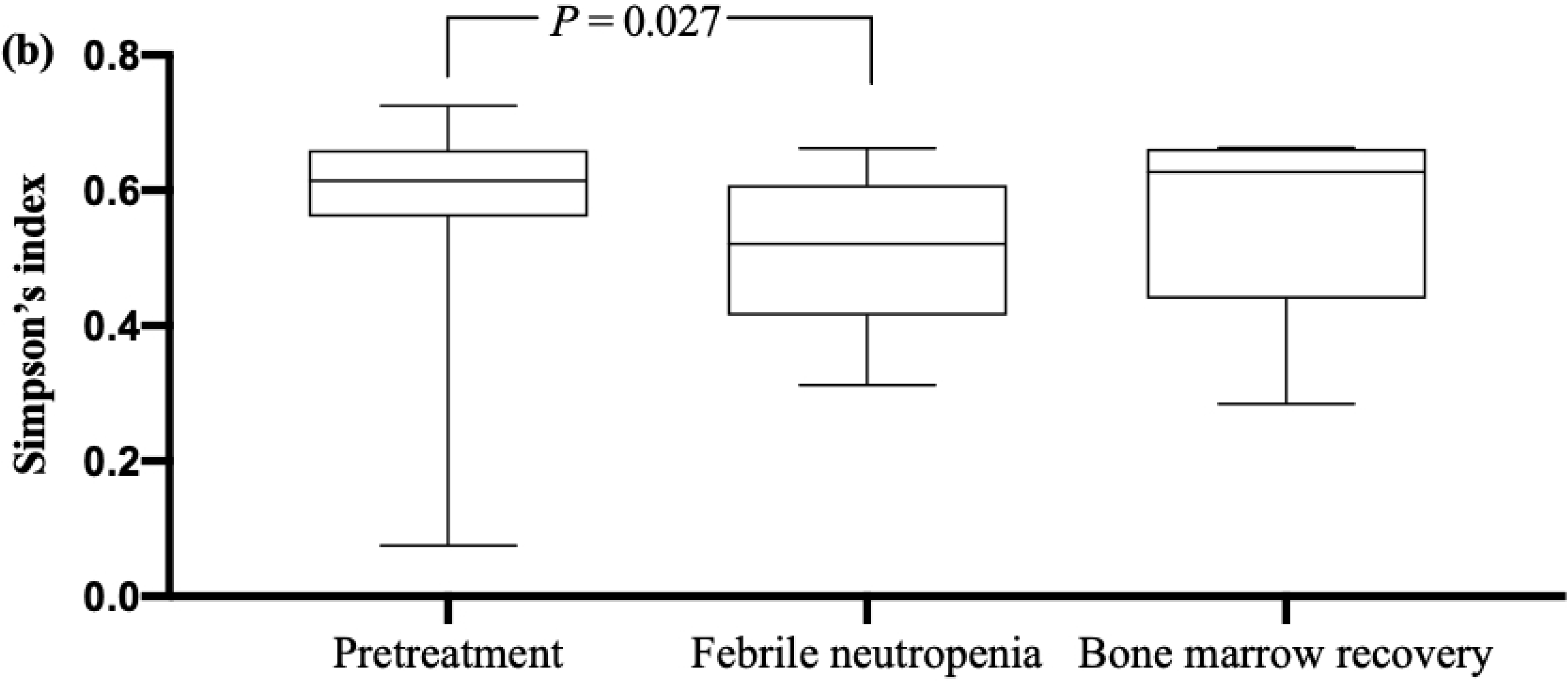
Diversity indexes at phylum level. (a) Shannon’s index values; (b) Simpson’s index values comparison between group by one-way ANOVA (comparisons between all groups) and Wilcoxon’s signed-rank test (comparisons between the paired samples). The 25th and 75th percentile are shown in the box plot. The median is indicated by horizontal solid lines. The bars indicate the minimum and maximum values.

## Discussion

This study demonstrated the alteration of gut microbiota in newly diagnosed adult AML patients who received first cycle of induction chemotherapy with carefully controlled factors that certainly affect composition of gut microbiota and entire participants developed first episode of NF. We hypothesize that even the first cycle of intensive chemotherapy, which able to contribute the NF, might reveal evidences of intestinal bacteria flora shifting because all of febrile neutropenia patients require empirical treatment with broad spectrum antibiotics with might eradicate particular taxa that affect host immunity and chemotherapy capable to extensively damage the gut barrier including neutropenia itself may affect patient’s intestinal bacteria.[28]

Although the majority of previous reports of gut microbiota were predominance of the patients who received the stem cell transplantation that neutropenic period might be intense and longer duration of bone marrow recovery compare to other kinds of chemotherapeutic regimens. For the allogeneic stem cell transplant recipients; the increase of relative abundance of *Enterococcus* and *Proteobacteria* during peri-transplant period significantly associate to bloodstream infection (BSI) and likelihood of bacterial translocation [7][29], loss of diversity with domination by single taxa including exposure to particular anti-anaerobic antibiotics subsequently escalate the risk of graft-versus-host disease (GVHD) and mortality rate.[10, 11, 30] Moreover, there was a study of pretreatment gut microbiota to predict the chemotherapy-related blood stream infection and machine learning was used to create the BSI risk index scoring system for non-Hodgkin lymphoma patients who underwent autologous stem cell transplantation. [31] All of these results confirmed that intestinal tract microbial diversity play a major role on the outcomes of treatment and pursue to multiple complications.

This study found the significant loss of fecal microbial diversity during neutropenic period with domination of phylum Firmicutes and significant increment of *Enterococcus* at genus level. This finding was in agreement with the previous stool microbiota studies in adult AML patients. [6, 32, 33] The increase of bacterial abundance in Enterococcaceae and Streptococcacea family in the Firmicutes phylum was previously reported as the strong predictor of infectious complications in pediatric acute lymphoblastic leukemia (ALL) and adult AML patients. [34, 35]

The single large study of gut microbiota in AML during induction chemotherapy [36] demonstrated the longitudinal analysis of oral and stool microbiota measurement that could assist the mitigation of infectious complications. The significant decreasing in both oral and stool microbial diversity were observed over the course on induction chemotherapy with good correlation between both sites of sample collection. The patients who lost microbial diversity were significantly more likely to detect a microbiologically documented infection within 90 days after treatment. The comparison study of gut dysbiosis between intensive chemotherapy and allogeneic hematopoietic stem cell transplantation also supported the similar loss of microbial diversity and domination of low-diversity communities by *Enterococcus* during period of intense neutropenia. [6] Additionally, this study also noted to the prominent reduction of *Bacteroides* and *Escherichia* (subset of Bacteroidetes and Proteobacteria family, respectively) while NF period. However, there were inconclusive reports on the alterations of Bacteroidetes family with scarce clinical data correlation, Proteobacteria family was relatively lack of information on changing pattern with a single report as predictive risk of febrile neutropenia and all of these reports performed solely in pediatric ALL patients. [33, 35, 37]

Moreover, the ethnicity and geographic location are considered to influent the composition of the gut microbiota. Interindividual differences in the intestinal microbiota profiles were intensively explained by an individual’s host locations [38] and even the same environment but ethnics variations also showed significant difference of gut microbiota patterns. [39] Unfortunately, the gut microbiota studies with association of ethnics and geographic region were explored in healthy persons that not reflect to setting of patients with hematologic cancer. In particular, the present knowledge of how the composition of the gut microbiota relates to patient health is principally based on investigations of European and North American populations which may limit the generalizable properties of microbiome-based applications for personalized medicine.[40] This study was exclusively resulted of Asian population with uniform pattern of clinical course including the treatment-naïve patients, homogenous course of chemotherapy and no previously used of antimicrobial prophylaxis in all AML patients.

Our study has several limitations to be concerned. Firstly, the relatively small sample size of the study and missing data might preclude to interpret the results due to lack of statistical power. However, this obstacle could be eliminated by carefully design to strictly control of stool samples collection on treatment-naïve patients, without factors affecting the intestinal microbiota such as previously received antibiotics, probiotics and nasal tube feeding or parenteral nutrition during study period and careful selection of stool sampling at the first date of febrile neutropenia and the first date of bone marrow recovery. Absence of infectious complications and clinical data correlation among shifting of gut microbiota is one of the weakness affecting the practical application.

In conclusion, the first-time treated adult AML patients during NF expressed the loss of fecal microbial diversity with greater amount of Firmicutes, but reverse trend in Bacteroidetes and Proteobacteria. *Enterococcus* was predominated genus with decrease of *Bacteroides* and *Escherichia* were disclosed. These findings revealed the evidence for conducting further microbiota researches in adult AML patients and introducing the way to launch additional larger studies to confirm the valuable clinical decision as individualize patient for antimicrobial de-escalation and prediction of anticipated complications.

## Acknowledgements

This study was funded by the research grant from Faculty of Medicine, Chiang Mai University (Chiang Mai, Thailand) and Omics sciences and bioinformatics center of Chulalongkorn University (Bangkok, Thailand) for sequencing and obtaining the gut microbiota data.

## Author contributions

**Conceptualization:** Thanawat Rattanathammethee, Pimchanok Tuitemwong, Parameth Thiennimitr

**Data curation:** Thanawat Rattanathammethee, Pimchanok Tuitemwong, Parameth Thiennimitr

**Formal analysis:** Thanawat Rattanathammethee, Pimchanok Tuitemwong, Phinitphong Sarichai, Sarisa Na Pombejra

**Funding acquisition:** Thanawat Rattanathammethee, Pimchanok Tuitemwong

**Investigation:** Thanawat Rattanathammethee, Pimchanok Tuitemwong

**Methodology:** Thanawat Rattanathammethee, Pimchanok Tuitemwong, Parameth Thiennimitr, Phinitphong Sarichai, Sarisa Na Pombejra

**Writing - original draft:** Thanawat Rattanathammethee, Pimchanok Tuitemwong

**Writing - review & editing:** Parameth Thiennimitr, Pokpong Piriyakhuntorn, Sasinee Hantrakool, Chatree Chai-Adisaksopha, Ekarat Rattarittamrong, Adisak Tantiworawit, Lalita Norasetthada

